# Learning enhances sensory processing in mouse V1 before improving behavior

**DOI:** 10.1101/087130

**Authors:** Ovidiu Jurjut, Petya Georgieva, Laura Busse, Steffen Katzner

**Affiliations:** Werner Reichardt Centre for Integrative Neuroscience, University of Tübingen, 72076 Tübingen, Germany; Division of Neurobiology, Department Biology II, LMU Munich, 82151 Munich, Germany

## Abstract

A fundamental property of visual cortex is to enhance the representation of those stimuli that are relevant for behavior, but it remains poorly understood how such enhanced representations arise during learning. Using classical conditioning in mice, we show that orientation discrimination is learned in a sequence of distinct behavioral stages, in which animals first rely on stimulus appearance before exploiting its orientation to guide behavior. After confirming that orientation discrimination under classical conditioning requires primary visual cortex (V1), we measured, during learning, response properties of V1 neurons. Learning improved neural discriminability, sharpened orientation tuning and led to higher contrast sensitivity. Remarkably, these learning-related improvements in the V1 representation were fully expressed before successful orientation discrimination was evident in the animals’ behavior. We propose that V1 plays a key role early in discrimination learning to enhance behaviorally relevant sensory information.

## Introduction

How does sensory processing in visual cortex change during learning of a stimulus’ behavioral relevance? A fundamental property of neurons in visual cortex is to enhance the representation of those stimuli that are relevant for behavior (reviewed in Gilbert and Li, 2013; Gavornik and Bear, 2014; Maunsell, 2015). The behavioral relevance of any given stimulus needs to be learned, though, and little is known about how visual sensory processing changes, as the significance of that stimulus becomes clear. While long-lasting alterations in visual cortical processing after mere exposure to specific stimuli (Frenkel et al., 2006; Cooke and Bear, 2010; Cooke et al., 2015) or after learning (Crist et al., 2001; Schoups et al., 2001; Ghose et al., 2002; Schwartz et al., 2002; Furmanski et al., 2004; Li et al., 2004; Yang and Maunsell, 2004; Raiguel et al., 2006; Hua et al., 2010; Jehee et al., 2012; Goltstein et al., 2013) have been documented amply, demonstrations of dynamic changes during learning are rare (Law and Gold, 2008; Li et al., 2008; Poort et al., 2015).

How animals learn about the relevance of any stimulus has been extensively studied at the level of behavior, where much of the research has relied on classical conditioning. Classical conditioning has the advantage that the occurrence of each lesson, such as a pairing of a stimulus with a reward, is entirely under the experimenter’s, and not the animal’s control (Pearce and Bouton, 2001). Therefore, the speed of learning can be elegantly manipulated, and its time course precisely quantified (Balsam and Gallistel, 2009).

Classical conditioning is often viewed as a reflexive, automatic type of learning; it does, however, involve substantial cognitive processes related to attention and decision-making. Support for these processes comes from the analysis of single-subject learning curves, which reveal in many learning paradigms step-like changes in behavior. Such step-like changes are inconsistent with a gradual strengthening of an association (Rescorla and Wagner, 1972) but can readily be explained within the framework of perceptual-decision making (Gallistel and Gibbon, 2000): across trials, the animal accumulates evidence that a given stimulus reliably predicts an important event and decides to show the conditioned response once the evidence is strong enough. The implication of these findings is that learning curves of individual subjects can be decomposed into a sequence of well-defined stages.

Whether progress in behavior and changes in sensory processing occur in parallel, or follow different time courses, is a matter of debate. Changes in sensory processing have been extensively investigated in primate studies of perceptual learning (reviewed in Gilbert and Li, 2012). In those rare cases where neural activity was measured during task performance, improvements in behavioral sensitivity were largely paralleled by improvements in neural representations (Law and Gold, 2008; Li et al., 2008). Similar observations were recently made in mouse V1, where chronic 2-photon calcium imaging revealed changes in population selectivity in parallel with progress in discrimination learning (Poort et al., 2015). Other studies, however, have documented that learning-related changes in neural processing can be dissociated in time from behavioral effects. In those cases, learning-related changes in neural activity are rather transient; they are pronounced during initial phases of training but relax, with additional practice, to pre-training levels although behavioral performance remains high (Zelcer et al., 2006; Yotsumoto et al., 2008; Sarro et al., 2015).

Here, we exploited the advantages of classical conditioning and quantified, in individual mice, the time course of orientation discrimination learning. Learning indeed occurred in a sequence of distinct stages, which were marked by qualitative changes in behavior. During traversal of these stages, we measured response properties of V1 neurons and found that the neural representation of the stimuli was improved to full extent well before animals showed any behavioral sign of orientation discrimination. These neural signatures likely reflect a key role of V1, which might enhance, early in discrimination learning, behaviorally relevant visual information.

## Materials and methods

We used 16 mice (2-6 months old, 11 males and 5 females), 13 of the C57BL/6J wild type strain and 3 of the PV-Cre strain B6;129P2-Pvalbtm1(cre)Arbr/J (JAX stock number 008069). All procedures were carried out in compliance with the European Communities Council Directive 2010/63/EC and the German Law for Protection of Animals; they were approved by the local authorities following appropriate ethics review.

### Surgical protocol

Anesthesia was induced with Isoflurane (3%) and maintained throughout the surgery (1.5%). A custom lightweight aluminum head post was attached to the anterior part of the skull (OptiBond FL primer and adhesive, Kerr dental; Tetric EvoFlow dental cement, Ivoclar vivadent); two miniature screws (00-96 x 1/16 stainless steel screws, Bilaney) were implanted over the cerebellum serving as reference and ground for electrophysiological recordings. Before surgery, atropine (Atropinsulfat B. Braun, 0.3 mg/kg sc) and analgesics (Buprenorphine, 0.1 mg/kg sc) were administered, and eyes were protected with ointment (Bepanthen). The animal’s temperature was kept at 37° C via a feedback controlled heating pad (WPI). Antibiotics (Baytril, 5 mg/kg sc) and a longer lasting analgesic (Carprofen, 5 mg/kg sc) were administered for 3 days post-surgery. Expression of channelrhodopsin (ChR2) in PV-Cre mice was achieved by injecting into V1 of anesthetized animals, through a small craniotomy, the adeno-associated viral vector rAAV5.EF1a.DIO.hChR2(H134R)-EYFP.WPRE.hGH (Penn Vector Core, University of Pennsylvania). A Picospritzer III (Parker) was used to inject the virus at multiple depths while gradually retracting the pipette. Mice were given 7 days to recover before they were habituated to the experimental setup. Before electrophysiological recordings, a craniotomy (~ 1.5 mm^2^) was performed over V1, 3 mm lateral to the midline and 1.1 mm anterior to the transverse sinus (Wang et al., 2011). The craniotomy was sealed with Kwik-Cast (WPI), which was removed and re-applied before and after each recording session.

### Experimental setup and visual stimulation

Mice were put on an air-cushioned Styrofoam ball and head-fixed by clamping their head-post to a rod. Movements of the ball were recorded at 90 Hz by two optical mice connected to a microcontroller (Arduino Duemilanove). A computer-controlled syringe pump (Aladdin AL-1000, WPI) delivered precise amounts of water through a drinking spout, which was positioned in front of the animals’ snout. Attached to the spout was a piezo element, which registered licking behavior (Schwarz et al., 2010). The drinking spout was present only during the conditioning experiments, and removed during measurements of orientation tuning. Visual stimuli were generated with custom-written software (**<u>https://sites.google.com/a/nyu.edu/expo/home</u>**) and presented on a liquid crystal display (LCD) monitor 25 cm in front of the animals’ eyes (Samsung 2233RZ, mean luminance of 50 cd/m^2^, refresh rate 120 Hz). Luminance non-linearities of the display were corrected with an inverse gamma lookup table, which was regularly obtained by calibration with a photometer. Stimuli consisted of sinusoidal or square wave gratings, which were 40-50 deg in diameter, and positioned to maximally overlap with the receptive fields (RFs) of the recorded neurons. Temporal frequency was 1.5 Hz, spatial frequency 0.02-0.04 cycles/deg. The setup was enclosed with a black fabrics curtain. Eye movements were monitored under infrared illumination using a zoom lens (Navitar Zoom 6000) coupled to a camera (Guppy AVT, frame rate 50 Hz). Optical stimulation was delivered via an optical fiber coupled to a light-emitting diode (LED; Doric lenses) with a center wavelength of 473 nm, which was driven by and LED driver (LEDD1B, Thorlabs). The fiber was 1 mm^2^ in diameter and the LED light intensity, measured at the tip of the fiber, was 0.7-3.5 mW/mm^2^. Before every recording session the fiber was positioned over the craniotomy, perpendicular to the brain surface, at a distance of ~1 mm using a micromanipulator. The animal’s eyes were shielded from the blue light by a sheet of black non-reflecting aluminum foil placed around the stimulation site.

### Initial behavioral training

After recovery from the surgery, animals were placed on a water restriction schedule until their weight dropped below 85 % of their ad libitum body weight. During this time, mice were habituated to head-fixation on the ball and delivery of water through the spout, which was triggered by the animal’s licking in the absence of visual stimuli. The animals’ weight and fluid consumption were monitored and recorded on each day, and the animals were checked for potential signs of dehydration. After the weight had stabilized the classical conditioning sessions started. These were typically performed 7 days a week, and only during these sessions mice received water.

### Conditioning paradigm

Animals were trained to discriminate between 2 gratings, one of them drifting down and to the left (-45 deg), the other one down and to the right (45 deg). We chose these two directions because they provide comparable sensory drive to neurons preferring horizontal gratings (0 deg), which are prominent in mouse V1 (Kreile et al., 2011). Gratings could vary in contrast (6 levels: 1, 2, 4, 6, 16, 40 and 100%). On a given trial, contrast and direction of movement were determined pseudo-randomly, and the grating was presented for 3 s. The presentation of one grating (-45 deg) was always followed by reward delivery (2-7 μl); the other grating had no consequences. The intertrial interval was 8-15 s plus a random delay drawn from an exponential distribution with a mean of 10-15 s. With such an interval, the animals cannot predict the onset of the stimulus. A single session consisted of 120-240 trials per day, divided into blocks of 60 trials.

### Electrophysiological recordings

Extracellular recordings were performed with 32-channel linear silicon probes (Neuronexus, A1x32-5mm-25-177-A32). Electrodes were inserted perpendicular to the brain surface and lowered to ~900 μm below the surface. Wideband extracellular signals were digitized at 30 kHz (Blackrock microsystems) and analyzed using the NDManager software suite (Hazan et al., 2006). Local field potentials (LFPs) were extracted by downsampling (1250 Hz) and filtering (4-250 Hz bandpass) the wideband signal. To isolate single neurons from linear arrays, we grouped adjacent channels into 5 equally sized “virtual octrodes” (8 channels per group with 2 channels overlap). Using an automatic spike detection threshold (Quiroga et al., 2004), spikes were extracted from the high-pass filtered continuous signal for each group separately. The first 3 principal components of each channel were used for automatic clustering with KlustaKwik (K. D. Harris, <u>http://klusta-team.github.io/klustakwik</u>), which was followed by manual refinement of clusters (Hazan et al., 2006). In the analyses of neural data we only considered high-quality single unit activity, judged by the distinctiveness of the spike wave shape and cleanness of the refractory period in the autocorrelogram.

### Measurements of eye position

The details are described in (Erisken et al., 2014). Briefly, we detected the pupil by convolving acquired camera frames with a symmetric Gaussian filter and applied a user defined threshold to obtain a binary image. We then applied a morphological opening operation, identified the most circle-like object as the pupil, and fitted an ellipse to determine the position of its center. To identify and compensate for translations of the eye parallel to the image plane, we also determined the position of a landmark, which could be either the 1^st^ Purkinje image of the infrared light, or a user defined point near the tear duct in the medial corner of the eye (Wallace et al., 2013). We computed relative pupil displacements by subtracting, for each frame the landmark position from the pupil position. To convert pupil displacements to angular displacements, we assumed that the center of eye rotation was 1.041 mm behind the pupil (Stahl et al., 2000). We defined saccades as changes in eye position greater than 2 degrees. Considering that the average mouse saccade lasts around 50 ms (Sakatani and Isa, 2007), we detected saccades by taking the difference of mean eye position 60 ms before and after each time point.

### Measurements of response properties

Before each classical conditioning experiment we mapped receptive field properties and measured orientation tuning. (1) RFs were mapped with a sparse noise stimulus, consisting of 5-deg full-contrast black and white squares, which were flashed, on a gray background, for 150 ms at a random location in a virtual 12x12 grid. Responses were fitted with a 2-dimensional ellipse to determine RF center, separately for ON‐ and OFF-subfields (Liu et al., 2010). (2) Orientation tuning was measured by presenting sinusoidal gratings at 100% contrast moving in a randomly selected direction (12 levels) for a duration of 2 s. Intertrial interval was 0.5 seconds. A blank screen condition (mean luminance) was included to estimate spontaneous firing rate. Each direction was presented 20 times. Orientation tuning curves were fit with a sum of two Gaussians, whose peaks were separated by 180 deg:

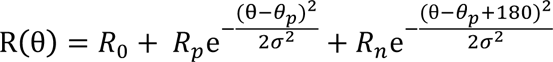

The function has 5 parameters: preferred orientation *θ_p_*, tuning width σ, baseline response *R_0_*, response at the preferred orientation *R_p_*, and response at the null orientation *R_n_*.

Contrast sensitivity was measured during the classical conditioning experiments, in which rewarded and unrewarded stimuli were presented at a range of contrasts. Contrast responses were fitted with the hyperbolic ratio function (Albrecht and Hamilton, 1982):

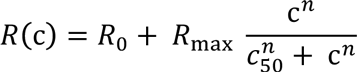

where c is stimulus contrast. The function has 4 parameters: baseline response *R_0_*, maximum response *R_max_*, semisaturation contrast *c_50_*, and exponent *n*. In a substantial fraction of V1 neurons responses did not saturate with increasing stimulus contrast. In these cases, estimates of *c_50_* hit the upper bound of meaningful values (i.e., 100%). To avoid this limitation, we took as a measure of contrast sensitivity the contrast, at which the neural response reached half of the maximum amplitude.

### Current source density analysis

We computed the current source density (CSD) from the second spatial derivative of the local field potential (Mitzdorf, 1985) in response to periodic visual stimulation. We smoothed the CSD in space using a triangular kernel (Nicholson and Freeman, 1975), and used a value of 0.4 S/m as measure of cortical conductivity (Logothetis et al., 2007) to approximate the CSD in units of nanoamperes per cubic millimeter. We assigned the contact closest to the earliest polarity inversion to the base of layer 4 (Schroeder et al., 1998). The remaining contacts were assigned to putative supragranular, granular and infragranular layers based on a cortical thickness of 1 mm and anatomical measurements of the relative thickness of individual layers in mouse V1 (Heumann et al., 1977).

### Analysis of visually evoked potentials

Visually evoked potentials (VEPs) were analyzed by computing the stimulus-triggered average from band-pass filtered (3-90 Hz) local field potentials recorded in layer 4 during measurements of orientation tuning. VEP amplitude was quantified by measuring trough to peak amplitude (Frenkel and Bear, 2004), where the trough was defined as the minimum value in the time interval from 0 to 100 ms after stimulus onset and the peak as the maximum value from 50 to 200 ms.

### Analysis of behavior

We quantified licking behavior on each trial by an index

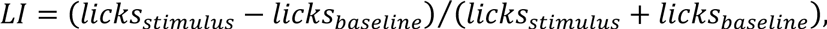

where *licks_stimulus_* is the number of licks during the final 1 second of stimulus presentation and *licks_baseline_* is the number of licks during the final 1 second before stimulus presentation. This index is bound between -1 and 1, with the two extremes indicating that licks exclusively occur during the baseline period or during grating presentation. We assessed learning progress by analyzing the cumulative sum of *LI*s (Papachristos and Gallistel, 2006), where changes in behavior become evident as changes in slope. Slope changes in the cumulative sum can more easily be seen than changes in raw indices across trials (Gallistel et al., 2004), and cumulative sums for all three learning stages can be compared in a single panel. To detect significant changes in slope, we performed the change the point analysis described in (Gallistel et al., 2001). To identify the transition from the naïve to the intermediate stage we examined *LI*s for the rewarded stimulus, and determined the trial, at which the first change point occurred. To identify the transition from the intermediate to the trained stage we performed the same analysis on the difference between the cumulative LIs for the rewarded and the unrewarded stimulus. To quantify the number of slope changes, we counted how many change points were needed before a post-change slope reached 80% of the terminal slope, which was based on the final half of the trials. For all but one analysis (28 out of 29), we used a statistical criterion of *p* < 10^−6^ for accepting changes in slope. In the remaining case, we lowered this criterion to *p* < 10^−5^. To validate the placement of change points, we performed ideal observer analyses (Macmillan and Creelman, 2005). For the first transition, we compared the distribution of lick rates compiled from the final second before stimulus onset against the distribution compiled from the final second during stimulus presentation. For this analysis we only used trials, in which the rewarded stimulus was shown and compared classification performance in the naïve versus later stages (i.e., intermediate and trained). For the second transition, we compiled distributions of *LI*s for rewarded vs. unrewarded trials and compared classification performance in the trained versus earlier stages (i.e., naïve and intermediate).

Animals were considered non-learners if their licking behavior failed to show a transition to the trained stage within 2000 trials. Three animals, however, did not receive the full amount of training, because decreasing quality of neural recordings led us to terminate learning sessions. Overall, of the 16 mice we used 1 dropped out before reaching the intermediate stage; 15 mice reached the intermediate stage and received further training. Of these 15 animals, 2 more dropped out by terminating learning sessions; 3 mice failed to reach the trained stage and were therefore classified as non-learners (n/a in Figure 2I).

To analyze running behavior, we transformed single-trial speed profiles within each session into binary vectors, in which 1s marked time points where the animal was running (speed > 1 cm/s) and 0s those where the animal was sitting (speed < 1 cm/s). Across trials we computed, for every point in time, the percentage of trials in which the animal was running, before averaging across sessions. To assess statistical significance, we computed, for each session, the average percentage of run trials within 0.1 to 1.5 s after stimulus onset, separately for each stimulus condition and learning stage. We then performed a mixed-design analysis of variance (ANOVA) involving the between-subjects factor learning stage (naïve vs. intermediate vs. trained) and the within-subject factor stimulus (rewarded vs. unrewarded).

#### Optogenetic suppression of V1 responses

After the animals had reached the trained stage, in which they could discriminate the two orientations, we combined electrophysiological recordings with photo-stimulation of parvalbumin-expressing (PV+) interneurons. In a random subset of trials (33%), we applied photo-stimulation concurrently with the presentation of the visual stimulus. In the experiments with optogenetic suppression we only used two levels of stimulus contrast (6% and 40%).

### Analysis of neural data

Individual animals typically contributed neural data to single learning stages.

In the analyses of neural discriminability and orientation selectivity we included neurons if (1) the sum of Gaussians explained at least 80% of the variance in responses, and (2) their trial-averaged firing rate during the presentation of their preferred stimulus was at least 1 spike/s.

Time courses of neural activity in Figure 5A,B are spike density functions computed by convolving single-trial spike trains with a Gaussian kernel (kernel resolution 10 ms, kernel width 100 ms) before averaging across trials.

#### Neural discriminability

We quantified how well individual neurons can discriminate the rewarded and unrewarded stimulus by extracting single-trial firing rates in a time window from 0.1 to 1.5 s after stimulus onset (excluding the transient part of the response) to compute a neural *d’*, defined as the difference in mean response to each stimulus, divided by the pooled standard deviation. During this window, mean lick rates were largely similar between stimuli across learning stages (rewarded vs. unrewarded stimulus; naïve stage: p = 0.97; intermediate stage: p = 0.43; trained stage: p = 0.07; interaction between rewarded vs. unrewarded stimulus and learning stage: p = 0.36, ANOVA). We sorted neurons according to preferred orientation into 4 bins centered on ‐ 45, 0, 45 and 90 deg, and performed an ANOVA on mean *d’* using the between-subject factors learning stage (naïve vs. intermediate vs. trained), layer (L2/3 vs. L4 vs. L5/6), and orientation bin. For post-hoc comparisons of individual means we used Tukey’s HSD test, with a 95% family-wise confidence level to correct for multiple testing.

#### Orientation selectivity

To compare counts of neurons across orientation bins and learning stages we performed a log-linear analysis of this multi-dimensional contingency table. To model the observed counts, we fitted a general linear model (GLM) with a Poisson link function considering the factors orientation bin (2 levels, -45 vs. 45 deg) and learning stage (3 levels). To compare the distributions of orientation preferences across learning stages without binning (Figure 5F) we used the multi-sample variant of the non-parametric Anderson-Darling test (Scholz and Stephens, 1987). This is an omnibus test, i.e. it provides a single test statistic to assess whether multiple distributions differ from each other. As an alternative, we compared pairs of distributions using the non-parametric Kolmogorov-Smirnov test, which led to the same conclusion of no difference between learning stages (all p > 0.44). To visualize how the laminar profile of tuning width σ was affected by learning (Figure 5H) we performed a non-parametric, locally-weighted, robust polynomial regression (lowess, Cleveland, 1979), using a parameter value of = 0.2 for the span of the smoothing window. To assess statistical significance, we ran an ANOVA on σ, using the same design as for *d’* (see *Neural discriminability*).

#### Contrast sensitivity

In the analysis of contrast sensitivity, we included neurons if (1) the hyperbolic ration function explained at least 90% of the variance in contrast responses, and (2) their maximum response was at least 1 spikes/s. Because contrast sensitivity is bound between 0 and 100, we assessed statistical significance of learning effects with the non-parametric Kruskal-Wallis omnibus test, followed by pairwise comparisons using the non-parametric Mann-Whitney test.

To assess whether locomotion affected contrast responses we analyzed data from a separate batch of animals, unrelated to the current study. For each neuron, we selected trials where the animals ran at least 80% and at most 20% of the time to compute contrast responses for locomotion and stationary periods. After fitting contrast responses with hyperbolic ratio functions (Albrecht and Hamilton, 1982) we compared the maximum response and the contrast at half the maximum response for locomotion versus stationary periods using paired t-tests.

#### Lick-triggered analysis of firing rates

We computed peri-event spike histograms centered on licks during a time window from 0.1 to 1.5 s after stimulus onset. We excluded from the analyses experiments with less than 10 licks and neurons with an average firing rate smaller than 1 Hz in a 200ms time interval centered around lick events. To compare distributions of lick-modulation indices across learning stages we used the multi-sample variant of the non-parametric Anderson-Darling test (see *Orientation selectivity*).

## Results

### Orientation discrimination learning unfolds as a sequence of distinct stages

We trained 16 mice, using classical conditioning, to discriminate the orientation of a grating stimulus and analyzed licking behavior to assess learning progress (Figure 1). The animals were head-fixed on a spherical treadmill in front of a computer monitor, on which we presented a drifting grating behind a stationary aperture. The gratings varied along 2 orthogonal orientations and 6 levels of contrast and were presented in a random order. The presentation of one orientation was immediately followed by a fluid reward, the other orientation had no consequences (Figure 1A,B). We measured licks as an indicator of learning progress, and observed pronounced changes in licking behavior across training sessions. Consider, for example, the sequence of training sessions shown in Figure 1C-E. In the ‘naïve’ stage, the animal licked spontaneously, but licking was unrelated to the visual stimulus (Figure 1C). In the ‘intermediate’ stage, the animal licked more frequently during the stimulus presentation; yet, the number of licks was similar for the rewarded and unrewarded orientation (Figure 1D). In this stage, the animal likely has associated the occurrence of either stimulus with the delivery of a reward. Finally, in the ‘trained’ stage, the animal showed anticipatory licks during the rewarded orientation, and largely suppressed licks during the unrewarded orientation (Figure 1E). In this stage, the animal has learned about the significance of the stimulus’ orientation.

**Figure 1.**
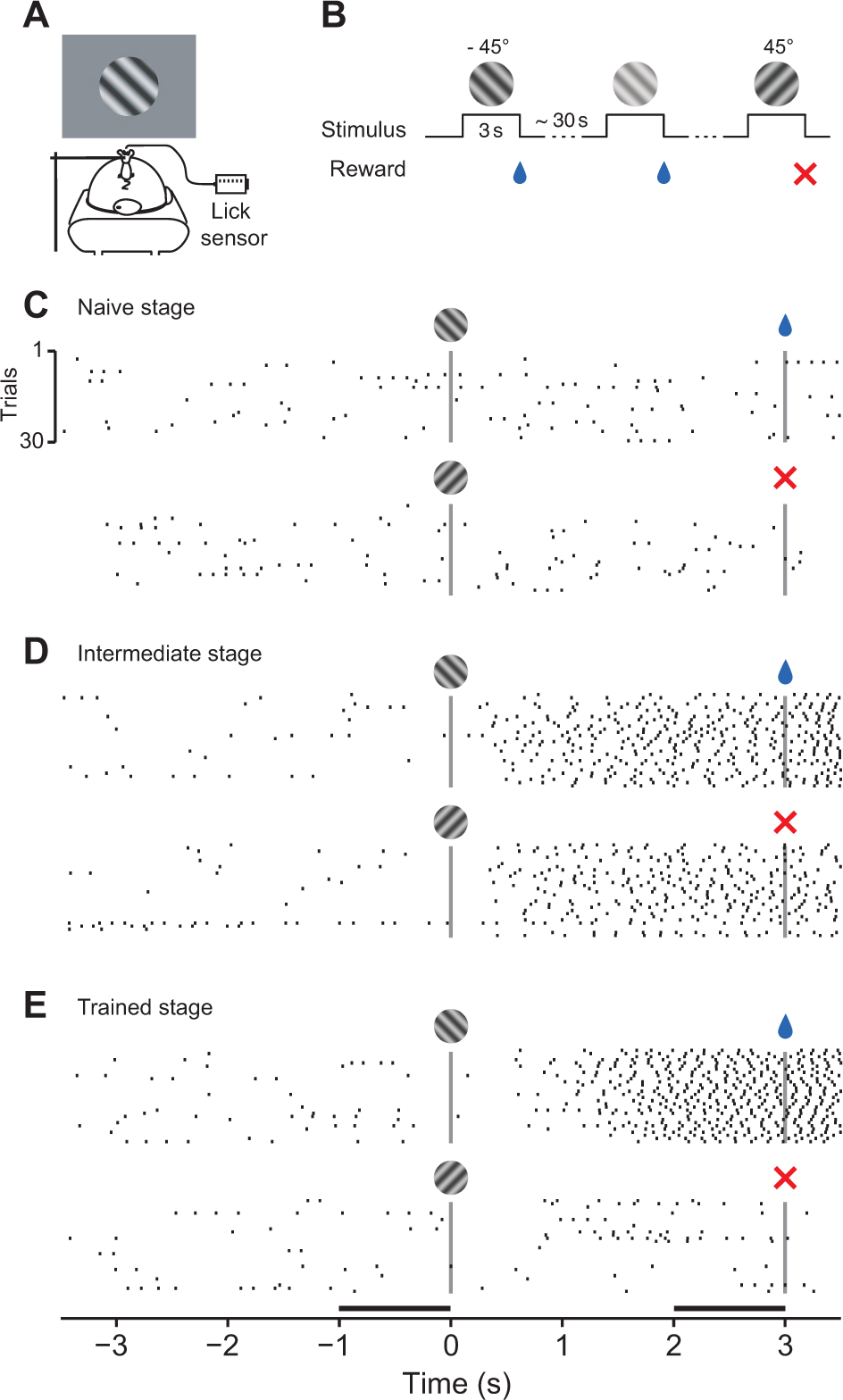
Classical conditioning of orientation discrimination. **(A)** Experimental setup. Head-fixed mouse on a spherical treadmill in front of a monitor showing oriented gratings. Reward was delivered through a drinking spout equipped with a lick sensor. **(B)** Discrimination learning paradigm. At irregular intervals, one of two oriented gratings was presented at a randomly selected contrast level. For one orientation (-45 deg), the presentation of the grating was immediately followed by a fluid reward; the other orientation (45 deg) had no consequences. **(C-E)** Licking behavior of one animal in 3 example sessions at different learning stages (mouse M22). Top: licks to the rewarded stimulus; bottom: licks to unrewarded stimulus; Vertical bars mark stimulus on‐ and offset. Horizontal bars in (E) mark the time windows used to quantify the strength of anticipatory licking.

We analyzed licking behavior of individual animals across learning stages and found that orientation discrimination learning can be decomposed into a sequence of distinct stages (Figure 2). To quantify licking behavior, we computed, for each stimulus separately, the cumulative sum of a lick index (LI), defined as the difference in lick rates between the final second during and the final second before stimulus presentation (black horizontal bars in Figure 1E), divided by their sum. Changes in the slope of this cumulative sum indicate changes in the animals’ behavior. We illustrate the sequence of distinct learning stages in three example mice (Figure 2A-C). During the naïve stage, cumulative LIs for the rewarded (blue) and unrewarded (red) stimulus fluctuate around zero. During the intermediate stage, the cumulative LIs rise with similar slopes because the animals increased lick rates after stimulus onset, irrespective of its orientation. During the trained stage, the cumulative LIs finally diverge, because the animals increase further anticipatory licking for the rewarded orientation, and/or suppress it for the unrewarded orientation. This point of divergence can best be seen as a change of slope in the difference between the cumulative LIs (Figure 2F-H). We used change point analyses (Gallistel et al., 2001) to find significant changes in slope and identify transitions between learning stages. We defined the transition from the naïve to the intermediate stage as the first change point in the cumulative LI for the rewarded stimulus (earliest blue circles in Figure 2A-C). Analogously, we defined the transition from the intermediate to the trained stage as the first change point in the difference of the two cumulative LIs (earliest black circles in Figure 2F-H). Relying on the very first change points is a conservative strategy and avoids contamination by trials from the subsequent stage. To quantify how distinct these transitions were, we determined how many change points were needed to reach 80% of the terminal slope, estimated from the final half of the trials.

**Figure 2.**
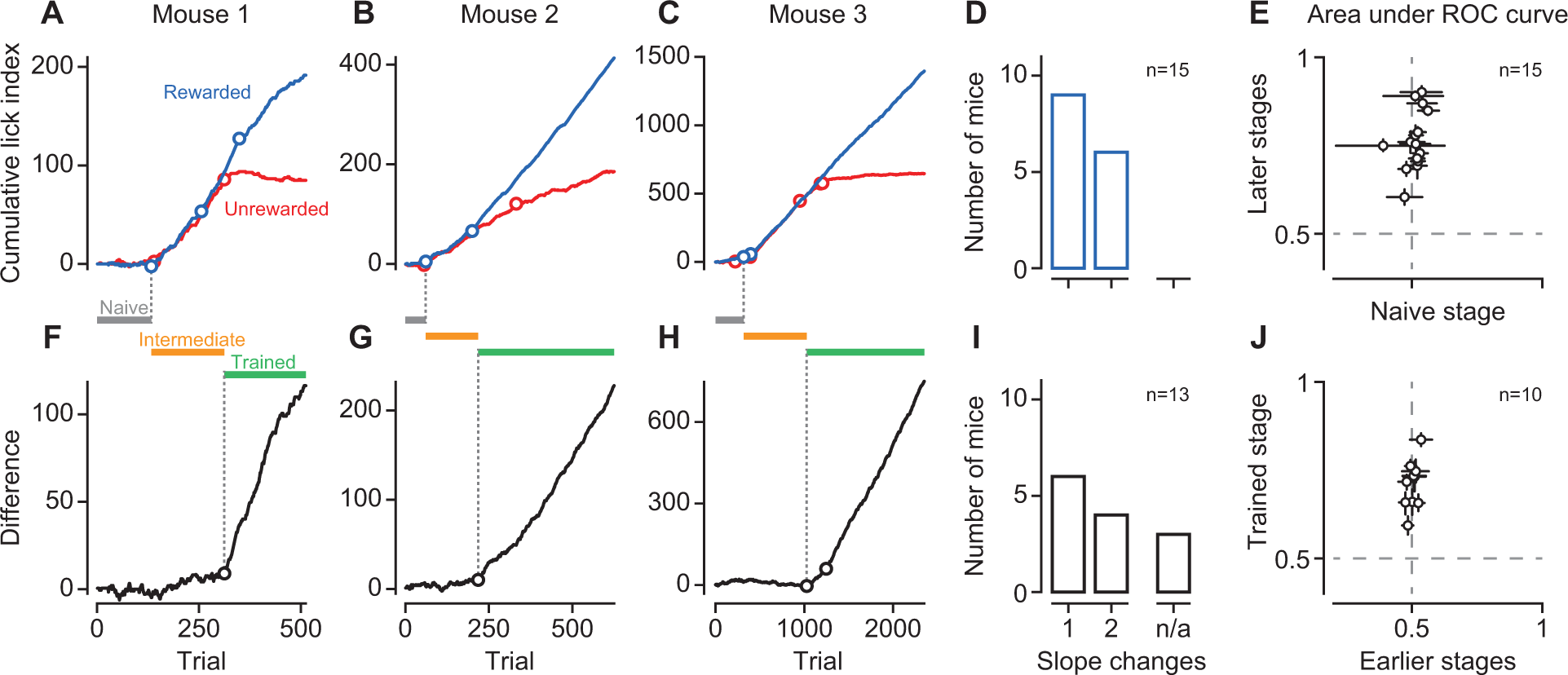
Orientation discrimination learning is characterized by a sequence of distinct stages. **(A-C)** Cumulative lick indices (LIs) across trials to the rewarded (blue) and the unrewarded (red) stimulus for 3 example mice (M22, M28, M42). Circles indicate significant change points. In (C), the second to last slope change for the unrewarded stimulus is concealed by the large number of trials, but becomes evident in the difference (H). **(D)** Distribution of the number of change points for the rewarded stimulus across mice (n = 15). **(E)** Ideal-observer analysis decoding stimulus presence from lick rates in naïve versus later (intermediate and trained) stages. **(F-H)** Same as (A-C), difference in cumulative LIs for rewarded and unrewarded orientations. **(I)** Same as (D), for the difference in cumulative LIs across mice, n/a gives the number of animals that failed to learn. **(J)** Ideal-observer analysis decoding stimulus orientation from LIs in early (naïve and intermediate) versus trained stages. Data points represent individual mice, error bars are 95%-confidence intervals.

All three example animals reached this criterion, in the cumulative LIs and in their difference, with only one or two changes in slope. These data indicate that there were few but clear-cut changes in behavior; before and after these changes behavior was largely constant for a stretch of trials.

This distinct sequence of learning stages was representative for the entire sample of animals (Figure 2D,I). For cumulative LIs to the rewarded orientation, all animals showed only one or two change points, after which performance reached the terminal slope (Figure 2D). Similarly, for the difference in cumulative LIs, all animals reaching the trained stage required at most two changes in slope (Figure 2I).

To further quantify licking behavior and validate the placement of change points, we performed an ideal observer analyses (Figure 2E,J). We separated trials based on change point locations and quantified how well an ideal observer could decode, from licking behavior, the presence of the stimulus or its orientation. In the naïve stage, licking was not predictive of stimulus presence (Figure 2E, lick rate before vs. during the rewarded stimulus, area under the receiver operating characteristic (AUROC): 0.51 ± 0.01, mean, standard error of the mean, n = 15 mice). Stimulus presence, however, could be well decoded from licking behavior during the final two stages (0.76 ± 0.02). Similarly, during the initial two stages, licking behavior was not predictive of stimulus orientation (Figure 2J, LIs for rewarded vs. unrewarded stimulus, AUROC: 0.50 ± 0.01, n = 10 mice); in the trained stage, however, stimulus orientation could be well predicted by licking behavior (0.71 ± 0.02).

Analyzing licking behavior separately for each level of stimulus contrast revealed that mice could detect the stimulus and discriminate its orientation at contrasts as low as 1%, and that stimulus contrast controlled the speed of learning (Figure 3). In all mice that were trained at multiple contrasts and traversed all learning stages (n = 7), licking behavior could be used to decode the presence of the stimulus even at 1% contrast (AUROC: 0.70 ± 0.04, n = 7, Figure 3A-D). Furthermore, for the majority of mice licking behavior reliably indicated, even at 1% contrast, stimulus orientation (0.62 ± 0.04, n = 7, Figure 3F-I). To examine how contrast affected the speed of learning we extracted the first change point at each contrast level and ranked them based on when they occurred; the later the change point, the higher the rank. The transition from naïve to intermediate, where the animals merely had to detect the presence of a stimulus, typically occurred faster with high-contrast stimuli (Figure 3E): Data points at 1% contrast cluster around 6, indicating that LIs at 1% contrast are last to change their slope. The transition from intermediate to trained, however, where the animals had to discriminate orientation, tended to occur faster for stimuli of intermediate contrasts (Figure 3J). In mice, improved visual performance at intermediate levels of stimulus contrasts has been reported earlier (Long et al., 2015). One potential explanation could be that our relatively large stimulus, at high levels of contrast, engages suppressive mechanisms outside the classical receptive field, which might impair decoding of stimulus orientation.

**Figure 3.**
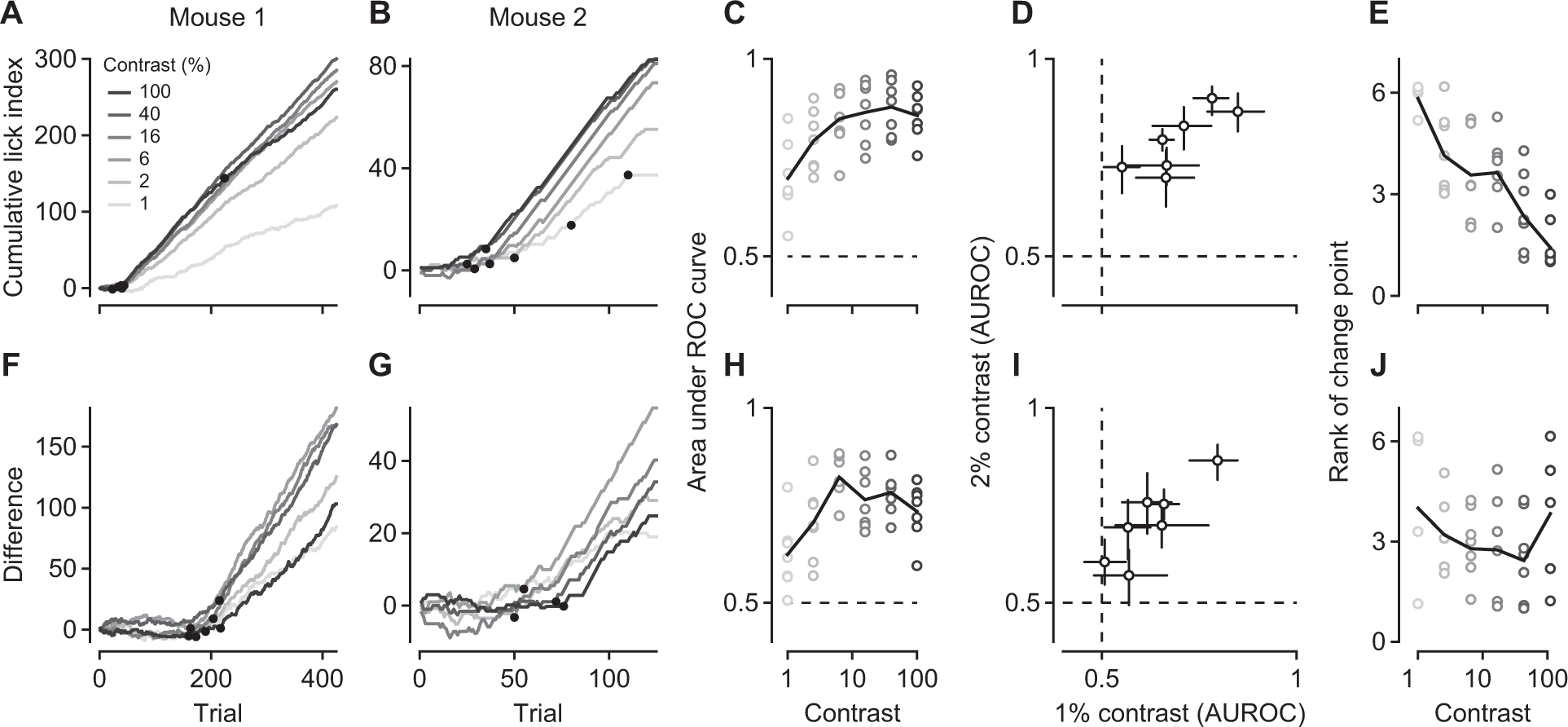
Orientation discrimination performance at different levels of stimulus contrast. **(A-B)** Cumulative LIs in response to the rewarded stimulus at different contrast levels for 2 example mice (M110, M42). Black dots indicate significant change points. **(C)** Area under the ROC curve as a function of stimulus contrast. At each contrast level, stimulus presence is decoded from lick rates in naïve versus late stages (intermediate and trained). Circles represent individual animals, black line is the mean across animals. **(D)** Same analysis as in (C), but showing contrast levels of 1 versus 2% only. Error bars represent 95% confidence intervals. **(E)** Rank of the first change point in the cumulative LIs as a function of stimulus contrast. Black line is the mean rank across animals. **(F-G)** Same as (A-B), but for difference in cumulative LIs to rewarded and unrewarded stimuli. **(H)** Same as (C), decoding stimulus orientation from LIs in naïve and versus late stages (intermediate and trained). **(I)** Same as (D), for data in (H). **(J)** Same as (E) for change points extracted from the difference in LIs. N = 7 mice.

Based on these analyses of behavior we conclude that orientation discrimination learning under classical conditioning is best described as a sequence of distinct stages, where transitions between stages are clear-cut and performance within stages largely constant. We next tested the hypothesis that traversing through these well-defined stages of learning was paralleled by changes in sensory processing in primary visual cortex.

Before studying how such distinct learning stages are reflected in cortical sensory representations, we confirmed that orientation discrimination in a classical conditioning paradigm requires V1. We found that discriminating grating orientation relies on V1 activity, because transient optogenetic suppression of V1 neurons impairs behavior (Figure 4). We expressed channelrhodopsin-2 (ChR2) in parvalbumin-positive (PV+) inhibitory interneurons, which we then stimulated with blue light to transiently reduce responses of V1 neurons. We performed current source density analysis (Mitzdorf, 1985) on the local field potentials to estimate the base of layer 4, and assigned neurons to putative supragranular (L2/3), granular (L4) and infragranular layers (L5/6) (Figure 4A). We photo-stimulated PV+ neurons, in a subset of randomly interleaved trials, during presentation of the stimuli at 40% or 6% contrast and observed profound reductions in responsiveness across the depth of cortex (Figure 4C-E). These reductions were stronger for stimuli at 6% contrast (mean suppression of 64 ± 4.7%) than for those at 40% contrast (52 ± 4.9%, p = 0.0006, paired t-test, n = 84 neurons). When reducing the responsiveness of V1 neurons, anticipatory licking during the reward-predicting stimulus was much weaker (Figure 4F,G, left, dashed blue line) compared to the control condition without photo-stimulation (solid blue line). Photo-stimulation did not seem to affect suppression of licking during the unrewarded stimulus (red lines), indicating that the light by itself does not lead to spurious or indifferent licking behavior. These effects of photo-stimulation not only were specific for the rewarded stimulus, but also depended on its contrast (Figure 4F,G, right). A constant intensity of photo-stimulation decreased anticipatory licking during the rewarded stimulus when presented at high contrast (black solid vs. dashed line), yet nearly abolished it when presented at low contrast (gray solid vs. dashed line). These findings show that our transient reduction of V1 activity interferes with the sensory processing of the stimulus, and does not simply cause random behavior in a disoriented mouse. These findings are consistent with previous work showing that mouse V1 is required for orientation discrimination (Glickfeld et al., 2013; Poort et al., 2015); we conclude that orientation discrimination under classical conditioning also relies on activity in V1.

**Figure 4.**
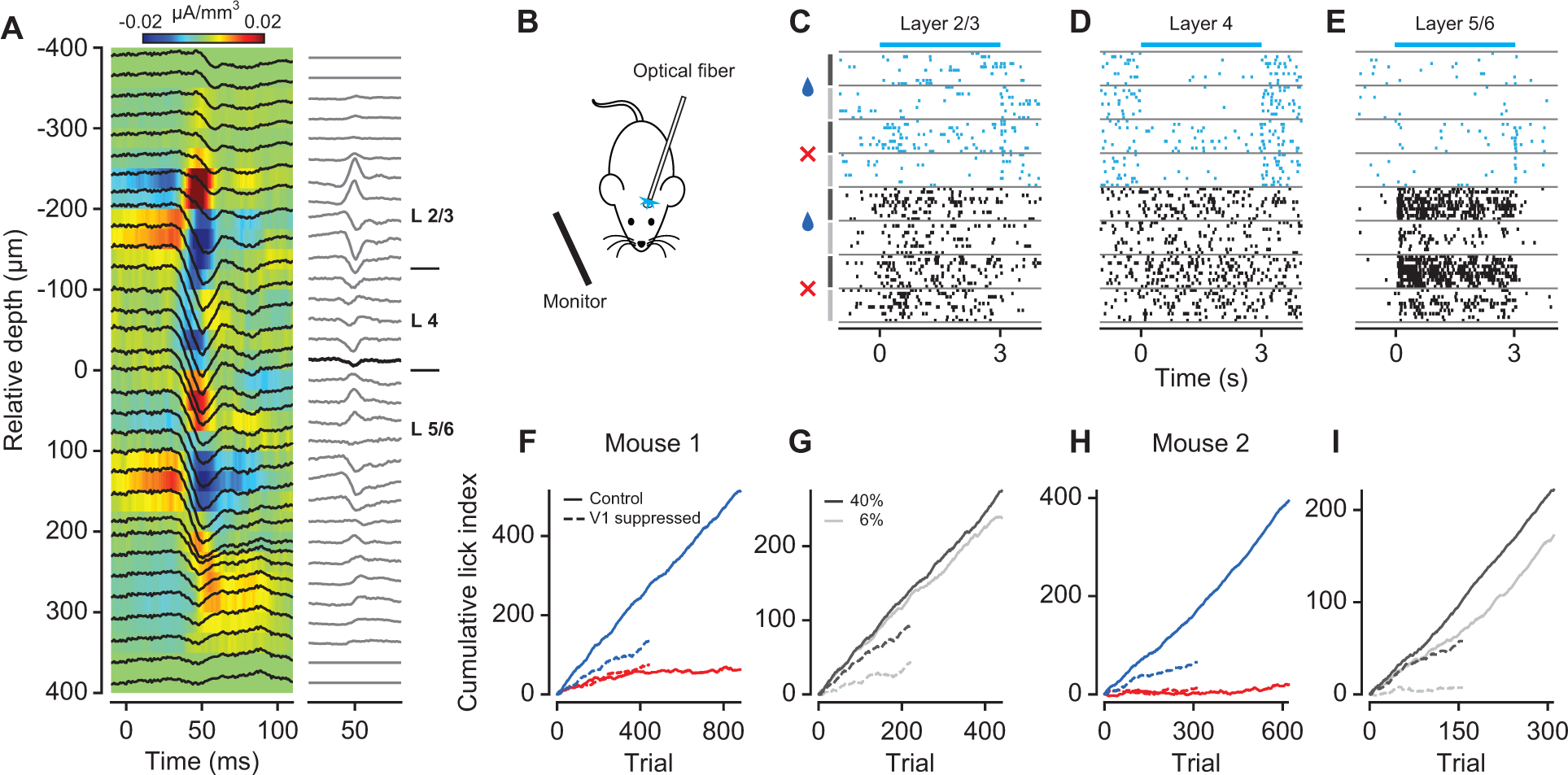
Transient optogenetic suppression of V1 neurons impairs behavioral performance. **(A)** left: local field potential (LFP, black traces) superimposed on the current source density (CSD) profile.: CSD traces, the bold line represents the base of layer 4. **(B)** V1 activity was suppresRightsed by unilateral photo-simulation of PV+ inhibitory interneurons expressing ChR2. **(C-E)** Raster plots of 3 example neurons from different V1 layers (units M120-33-8, -28, -47). Cyan: V1 suppression by PV+ photo-activation, Black: control condition. Stimulus conditions are separated by gray horizontal lines. Symbols and vertical lines on the left indicate stimulus orientation (blue drop: rewarded, red cross: unrewarded) and contrast (black: 40%, gray: 6%). **(F)** Cumulative lick index for one mouse (M117). Left panel: behavioral performance for rewarded (dark blue) vs. unrewarded (red) orientation during V1 suppression (dashed lines) and in control condition (solid lines). Right panel: rewarded orientation only, at 40% (black) and 6% (gray) contrast. **(G)** Same, for another mouse (M199).

### Learning shapes V1 responses well before the animals discriminate the stimuli

Having established that orientation discrimination in our classical conditioning paradigm relies on activity in V1 we recorded, during conditioning sessions, extracellular activity from ensembles of individual V1 neurons simultaneously across the depth of cortex. Within individual recording sessions, we interleaved conditioning experiments (‘the task’), with measurements of orientation tuning curves outside the context of this task.

To our surprise, we found that V1 neurons showed improved discriminability for the behaviorally relevant orientations already in the intermediate stage, where the animals still did not discriminate the stimuli (Figure 5). To quantify how well individual neurons could discriminate between the rewarded and unrewarded orientation (‘the relevant orientations’), we computed *d’*, defined as the difference in mean firing rates relative to the pooled standard deviation. The value of *d’* depends on the preferred orientation of a neuron: A neuron with preferred orientation close to the rewarded orientation, such as the example in Figure 5A, should have a positive d’; neurons, whose preferred orientation is somewhat in between, such as the example in Figure 5B, receive comparable drive from both orientations and their *d’* should be close to 0. To compute *d’*, we focused on a response time window from 0.1 to 1.5 s after stimulus onset (black horizontal bar in Figure 5A,B), where anticipatory licking behavior was largely similar between the two orientations (see Figure 1E).

**Figure 5.**
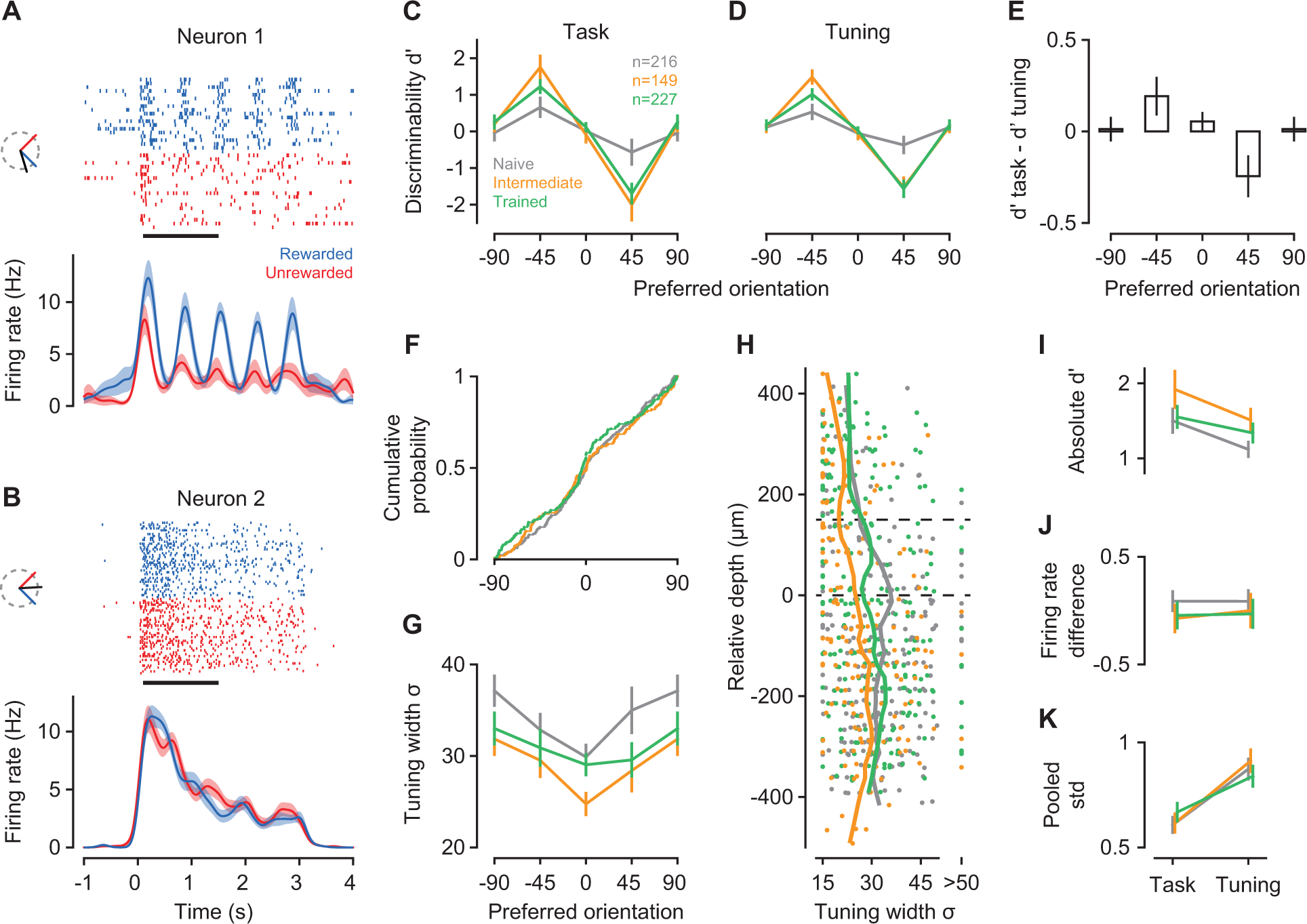
V1 neurons show improved discriminability and sharper orientation tuning already in the intermediate learning stage. **(A-B)** Spike rasters and density functions in response to the rewarded (-45 deg, blue) and unrewarded (45 deg, red) stimulus for two example neurons (units M22-14-16, M81-3-31). Insets illustrate preferred orientation (black) relative to rewarded and unrewarded orientation. **(C)** Neural discriminability (*d’)* during task performance in the naïve (gray), intermediate (orange) and trained stage (green) for neurons binned according to preferred orientation. **(D)** *d’*s computed from responses to the same stimuli during orientation tuning measurements. **(E)** Mean pair-wise differences in *d’* between task and tuning measurements across orientation bins. **(F)** Cumulative distribution of orientation preferences across learning stages. **(G)** Learning-related changes in tuning width as a function of orientation preference bin. **(H)** Tuning width (σ) and laminar location of all neurons recorded during the naïve, intermediate, and trained stage. Trends (vertical lines) were computed with locally weighted, robust regression (lowess). **(I)** Absolute *d’*s during task performance versus measurements of orientation tuning, separately for each learning stage. Included are only neurons from bins centered on -45 or 45 in (C). **(J)** same, for differences in firing rates between rewarded and unrewarded stimuli. **(K)** same, for pooled standard deviation. Measures in (J,K) are normalized by mean firing rate; n = 11 mice; numbers of neurons per learning stage are given in C; error bars represent ± 1 standard error of the mean.

We first examined V1 responses during task performance, and found that the improvements in discriminability were largely restricted to neurons with preferred orientations close to either of the relevant stimulus orientations (Figure 5C). Binning neurons by their orientation preference revealed that learning improved *d’*s in the two bins centered on the rewarded (-45) and unrewarded (45) orientation (interaction between learning stage and orientation bin, p < 10^−4^, ANOVA). Follow-up analyses confirmed that, for the rewarded stimulus, *d’*s were higher at the intermediate than at the naïve stage (p = 0.0065, Tukey’s HSD test); *d’*s measured during the trained stage, however, did not significantly differ from the other stages (p > 0.23 for both comparisons). Likewise, for the unrewarded stimulus, *d’*s were more negative during intermediate (p = 0.012) and trained stages (p = 0.03), compared to the naïve stage, but did not significantly differ from each other (p = 0.83). We tested whether this selective improvement of *d’*s would depend on laminar location, but found no evidence (interaction between learning stage, orientation bin, and layer, p = 0.43, ANOVA, data not shown).

To test whether improvements in neural discriminability require engagement in the task, we examined responses during measurements of orientation tuning; although we found improvements outside the context of the task, they were weaker than during task performance (Figure 5D,E). To evaluate neural discriminability without the animals being engaged in the task, we determined *d’s* from responses to the relevant orientations presented during measurements of orientation tuning, and also found significant improvements in *d’* with learning. In fact, the improvements during measurements of orientation tuning depended in the same way on the orientation preference of the neurons (Figure 5D, interaction between learning stage and orientation bin, p < 10^−4^, ANOVA). Although learning improved discriminability outside the task, *d’*s during orientation tuning were not as pronounced as during task performance (interaction between orientation bin and task, p = 0.0027). These additional improvements in discriminability during the task can best be seen by computing, separately for each neuron, the difference in *d’*s between task and tuning measurements (Figure 5E). During the task, the mean difference in *d’*s is positive for neurons preferring the rewarded stimulus (-45 deg, p = 0.027, one-sided t-test), negative for neurons preferring the unrewarded stimulus (45 deg, p = 0.014), and indistinguishable from zero for all other neurons.

Because the improvements in discriminability persisted outside the task, we analyzed the orientation tuning measurements to identify underlying neural mechanisms. The fraction of V1 neurons preferring either the rewarded or unrewarded stimulus was similar overall (main effect of bin: p = 0.65, log-linear analysis) and remained similar across learning stages (interaction between bin and learning stage: p = 0.74, data not shown). We also compared, without binning, the distributions of orientation preferences across learning stages and found them to be indistinguishable (Figure 5F, p = 0.30, Anderson-Darling test). These results indicate that classical conditioning of orientation discrimination does not induce any systematic shift in the distributions of orientation preferences.

While the distributions of orientation preferences did not show systematic shifts with learning, we found substantial sharpening of orientation selectivity, which was strong already in the intermediate learning stage (Figure 5G-H). Consistent with previous reports (Niell and Stryker, 2008; Sun et al., 2016), average tuning width had a clear laminar profile: it was narrower in L2/3 than in L4 and L5/6 (Figure 5H, means of 25 vs. 33 vs. 34 deg, main effect of layer, p < 10^−4^, ANOVA). Tuning width, however, further changed with learning (main effect of learning stage: p = 0.0038): throughout the depth of cortex, learning sharpened tuning width, particularly in the intermediate stage (orange, intermediate vs. naïve: p = 0.0026, Tukey’s HSD test). In the trained stage (green), tuning width was broader and more similar to the naïve stage (trained vs. intermediate: p = 0.061, trained vs. naïve: p = 0.44). Unlike the improvements in discriminability, this sharpening of orientation tuning width did not depend on a neuron’s orientation preference, but occurred globally across the entire population (interaction between learning stage and orientation bin: p = 0.84, Figure 5G). Together, these results indicate that the learning-related improvements seen outside the task were largely mediated by a sharpening of tuning width, which was most pronounced in the intermediate stage.

Given the additional benefit of task engagement, we sought to identify the properties of the neural response that further improved discriminability during the task. We focused on those neurons that showed learning-related improvements (orientation preference bins at -45 and 45 deg in Figure 5C) and examined how absolute *d’* changed between tuning measurements and task performance. As implied by the analyses of pairwise differences (Figure 5E), *d’*s were higher during the task than during tuning measurements (Figure 5I, main effect of task: p < 10^−4^, ANOVA), independent of the learning stage (main effect of learning stage: p = 0.13; interaction between task and learning stage: p = 0.48). We decomposed *d’* into differences in firing rates (rewarded – unrewarded, Figure 5J) and pooled standard deviation (Figure 5K), both of them normalized to mean firing rate. Performing the task did not affect differences in firing rates (main effect of task: p = 0.47), but strongly reduced firing rate variability (main effect of task: p < 10^−4^). Because these task-related improvements were present already in the naïve stage, they likely result from different levels of engagement. During tuning measurements, the animals passively view a sequence of oriented gratings; during the task they learn to focus on visual information to anticipate reward. These analyses indicate that being engaged in a perceptual task, as simple as learning to anticipate reward, can improve the reliability of sensory responses in V1.

We finally asked whether progress in orientation discrimination would affect processing of stimulus features other than orientation; to address this question we examined responses of V1 neurons to different levels of stimulus contrast (Figure 6). Because our rewarded and unrewarded stimulus had different orientations, comparing the contrast responses to these two stimuli within individual neurons would be confounded by differences in sensory drive (Figure 6A-C). We therefore compared contrast sensitivity in the population of V1 neurons, measured with the same stimulus, across the different learning stages. We fitted contrast responses with a hyperbolic ration function (Albrecht and Hamilton, 1982), and took as a measure of contrast sensitivity the contrast at half the maximum response (dotted lines in Figure 6A).

**Figure 6.**
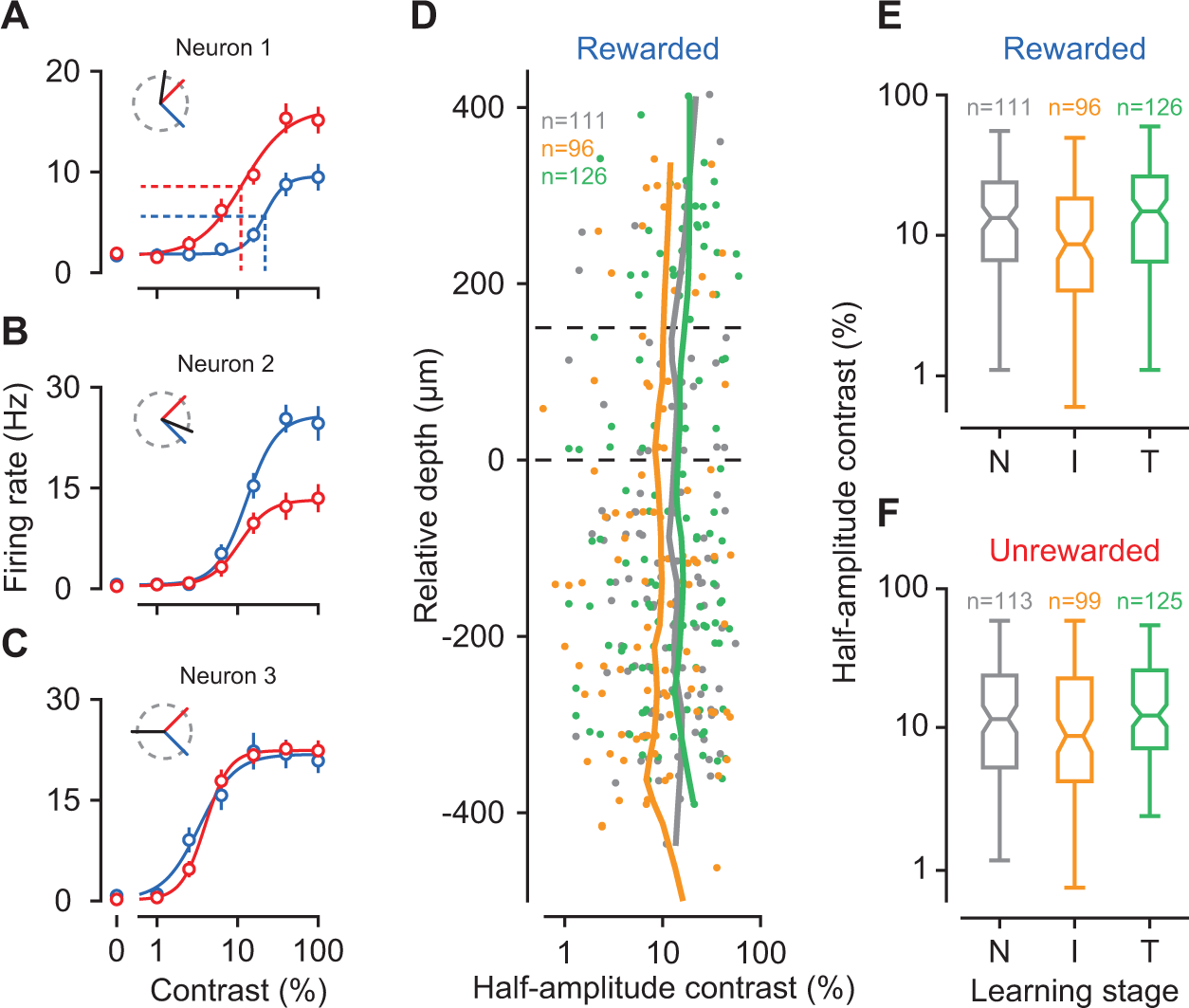
Contrast sensitivity in the population of V1 neurons. **(A-C)** Contrast responses of three example V1 neurons (units M110-30-5, M22-14-46, M154-6-15) for the rewarded (blue) and unrewarded (red) stimulus. Since the two stimuli are not identical, these contrast responses are strongly determined by the neuron’s preferred orientation (shown by black line in inset). Data are means ± 1 SEM. Dotted lines in (A) indicate contrast sensitivity, defined as the contrast at half the maximum response **(D)** Contrast sensitivity for the rewarded stimulus across the depth of cortex, separately for each learning stage. **(E)** Summary statistics of contrast sensitivity for the rewarded stimulus: Center marks are medians, box edges are the 25^th^ and 75^th^ percentiles, error bars give the range. **(F)** same, for unrewarded stimulus. N = 11 mice. Conventions as in Figure 5.

We examined responses to the rewarded stimulus, and found that contrast sensitivities were highest, across the depth of cortex, in the intermediate stage (Figure 6D). Pooling neurons across layers we found that contrast sensitivities differed between learning stages (Figure 6E, main effect of stage: p = 0.027, Kruskal-Wallis ANOVA). Follow-up comparisons revealed that contrast sensitivities were higher during the intermediate than during the naïve (p = 0.040, Mann-Whitney Test) and trained stage (p = 0.012); naïve and trained stages did not differ from each other (p = 0.45). For the unrewarded stimulus, the effects were less robust but gave similar results (Figure 6F, main effect of stage: p = 0.036, Kruskal-Wallis ANOVA). These data show that learning to discriminate orientations not only sharpens selectivity for orientation but also improves, in the intermediate stage, sensitivity for contrast.

Since our animals could rest or run at any time, we asked whether any of the observed changes in V1 processing could be related to the animals’ running behavior; we found that this was not the case (Figure 7). Because locomotion can increase the gain of sensory responses in mouse V1 (Niell and Stryker, 2010; Bennett et al., 2013; Polack et al., 2013; Erisken et al., 2014; Fu et al., 2014; Lee et al., 2014) and reduce trial-to-trial variability of membrane potential fluctuations (Bennett et al., 2013), it could, in principle, also bring about higher values of *d’* (Figure 5C-E). The increases in d’ we observed, however, were not confounded by differences in locomotion. We analyzed run-speed profiles from single trials (Figure 7A) to determine, for each point in time, the percentage of trials during which the animal was running. As the animals learned the task, locomotion locked to the structure of the trial (Figure 7B-D). During the window we used to analyze neural responses, however, the percentages of trials the animals spent running remained largely constant. Only in the trained stage, animals ran more in response to the rewarded than the unrewarded stimulus (Figure 7E, interaction between learning stage and reward: p < 10^−4^; ANOVA). Furthermore, *d’*s increased even outside the context of the task, where running behavior remained indistinguishable across the entire trial duration and all learning stages (Figure 7F-I, main effect of stage: p = 0.48, ANOVA). Similarly, we observed a narrowing of tuning width with learning outside the context of the task (Figure 5F-G), where locomotion was identical across learning stages. Furthermore, based on previous findings (Erisken et al., 2014; Lee et al., 2014), locomotion should not alter contrast sensitivity. We tested this directly by analyzing data obtained from a separate batch of naïve mice, unrelated to the current study. We compared contrast responses between locomotion and stationary trials and found that locomotion increased the maximum response (Figure 7J, p = 0.0003, paired t-test) without affecting contrast sensitivity (Figure 7K, p = 0.20). Together, these results argue against the possibility that the learning-related improvements in V1 sensory processing can be explained by the animals’ running behavior.

**Figure 7.**
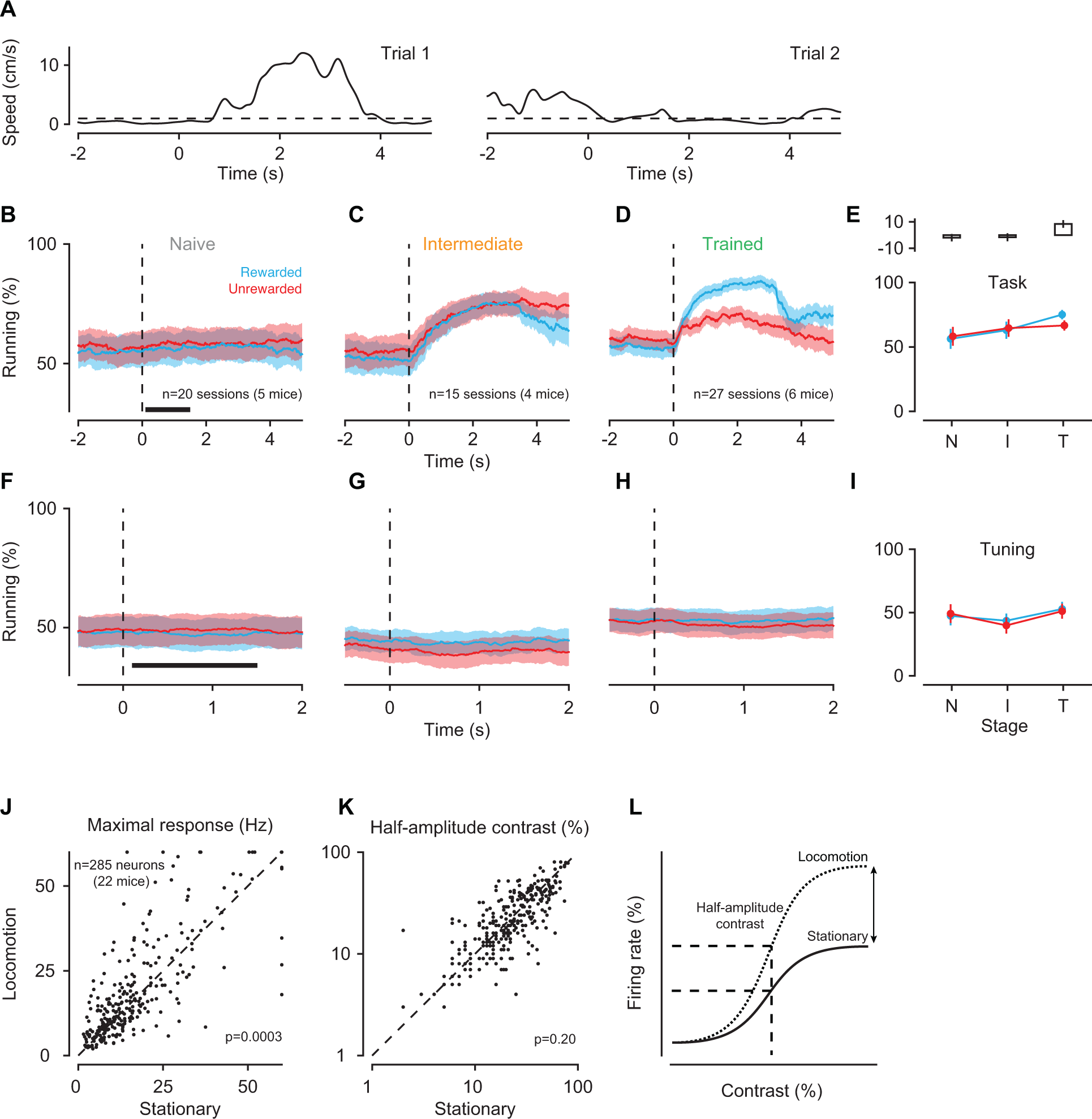
Running behavior during orientation discrimination learning. **(A)** Two single-trial speed traces (mouse M36). Dashed line is the speed threshold (1 cm/s) used to determine whether the animal was running or not. **(B)** Percentage of trials during which animals were running around the onset of the rewarded (blue) or unrewarded (red) stimulus in the naive stage. Shaded areas are ± 1 SEM across recording sessions. Dashed line indicates stimulus onset. Black bar marks the time window used for the analysis in (E). **(C)** Same, for intermediate stage **(D)** Same, for trained stage. **(E)** Bottom: Average percentage of run trials within 0.1 to 1.5 s after stimulus onset across learning stages (N = naïve, I = intermediate, T = trained). Top: mean pair-wise differences between rewarded and unrewarded conditions across recording sessions, together with their SEMs. **(F-H)** Same as (B-D), during measurements of orientation tuning but under identical sensory stimulation. **(I)** Same as **(E)**, during measurements of orientation tuning. **(J)** Maximum firing rate of V1 neurons during locomotion and stationary trials. **(K)** Contrast sensitivity of V1 neurons during locomotion and stationary trials. **(L)** Schematic summary of the effects: locomotion, on average, increases the maximum response without affecting contrast sensitivity.

These learning-related changes in visual processing also cannot be explained by artifacts of eye movements (Figure 8). We did observe occasional eye movements, which mainly occurred along the horizontal axis. Their overall frequency, however, was very low (naïve stage: 0.075 ± 0.04 Hz; intermediate stage: 0.096 ± 0.05 Hz; trained stage: 0.16 ± 0.07). Although mean saccade frequency changed with learning stage (main effect of stage, p = 0.0021, ANOVA), it was indistinguishable between the naïve and intermediate stage (p = 0.63, Tukey’s HSD test), where learning already shaped V1 responses. In addition, across learning stages, the distributions of eye positions were highly overlapping and centered on a default position.

**Figure 8.**
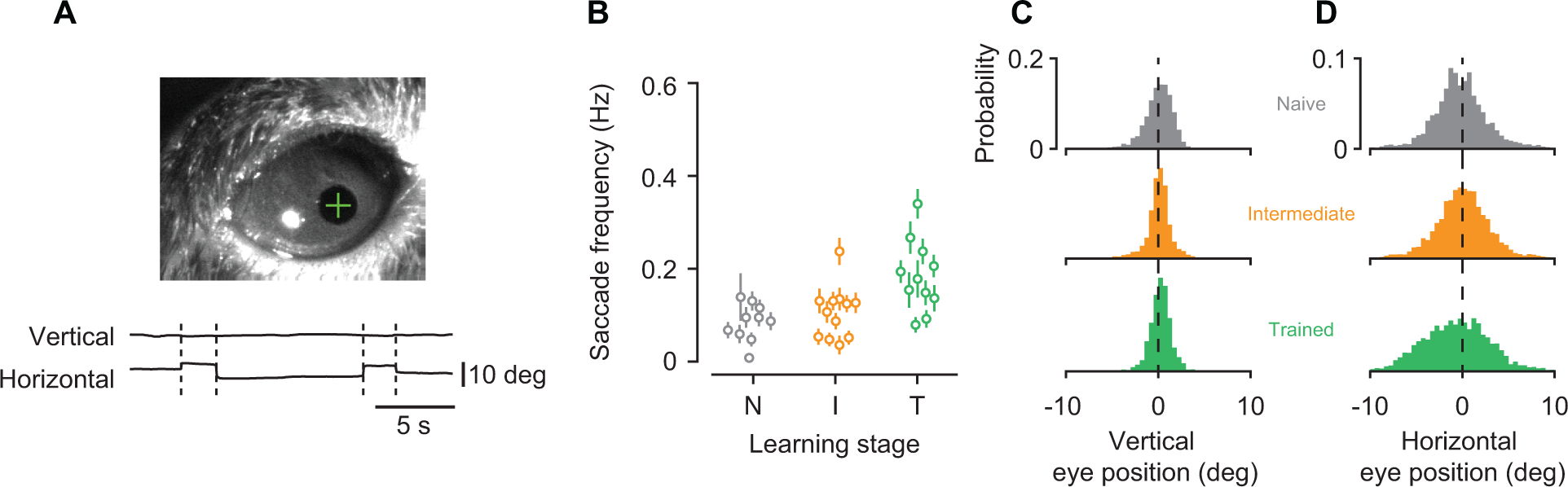
Learning-related changes in V1 processing were not an artifact of eye movements. **(A)** Eye position was measured by tracking the pupil under infrared illumination. Top: example image acquired by the eye-tracking camera. White spot is the cornea reflection of the infrared LED, green cross is the estimate of the pupil center. Bottom: example traces for vertical and horizontal eye position. Saccades are marked with dashed lines. **(B)** Saccade frequency within 0.1 to 1.5 s after stimulus onset across learning stages. Data points represent recording sessions, error bars indicate ± 1 SEM. **(C)** Distributions of vertical eye position. **(D)** Same, for horizontal eye position.

The learning-related changes in V1 sensory processing also cannot be explained by licking artifacts (Figure 9). First, we observed changes in d’ and tuning width outside the task, where the lick spout was completely removed from the setup. Second, we examined licking behavior during the task, and found that it only marginally influenced V1 responses. Within the window used to analyze the neural data, we constructed, for each neuron, peri-event spike histograms centered on licks (Figure 9A,B). We classified neurons as lick-modulated if the mean firing rate during a post-lick period of 100 ms was outside the 99-% confidence interval around the mean firing rate in the corresponding pre-lick period. The percentage of lick-modulated neurons was small (naïve stage: 3.5%, intermediate stage: 5.2%, trained stage: 3.5%), and statistically indistinguishable across learning stages (interaction between learning stage and lick-modulation: p = 0.10, log-linear analysis). We also computed a lick-modulation index, defined as the difference in mean pre‐ and post-lick firing rates divided by their sum, and compared the distributions of these indices across learning stages (Figure 9C). An omnibus test showed a significant difference between the three distributions (p = 0.023, Anderson-Darling test). Follow up-analyses then revealed that the indices were actually less extreme in the intermediate (orange) than in the naïve (gray, p = 0.0048) and trained stages (green, p = 0.057). These results argue against the possibility that the learning-related changes in V1 processing artificially arise from differences in licking behavior.

**Figure 9.**
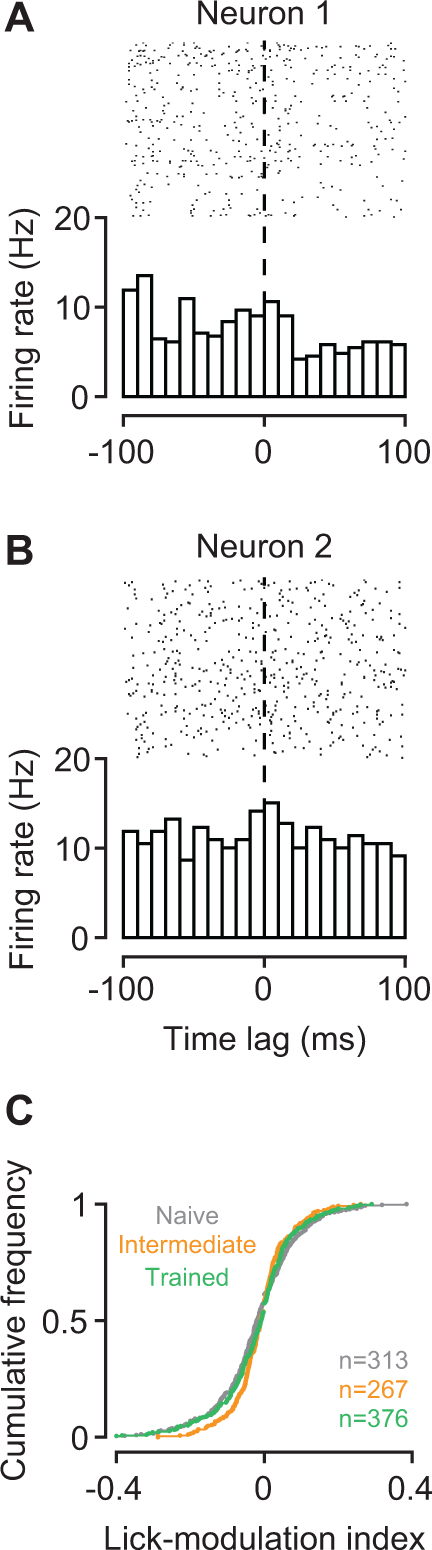
Learning-related changes in V1 processing are not an artifact of licking behavior. **(A,B)** Examples of peri-event spike histograms triggered on licks. Firing rates of neuron 1 are weakly reduced with licking; firing rates of neuron 2 are not affected (M53-1-8, M117-22-21). **(C)** Cumulative distributions of lick-modulation indices across the population of neurons during the naïve (gray), intermediate (orange) and trained stage (green).

Finally, we rejected the possibility that the learning-related changes result from a disproportionate exposure to the two orientations presented during the task (Figure 10). In mouse V1, repeated exposure to a certain orientation can lead to persistent enhancements of the visually evoked response to that stimulus (Stimulus-selective response potentiation (SRP), Frenkel et al., 2006; Cooke et al., 2015). SRP, however, is typically observed after prolonged presentation of the same stimulus at high levels of contrast (100-400 blocks of stimulus presentation at 100% contrast, (Frenkel et al., 2006)). We do not expect that our paradigm can lead to SRP. While we did present two orientations only during the task, 4 of our 6 levels of stimulus contrast were below what is required to produce SRP (> 12% contrast, Frenkel et al., 2006). At sufficiently high levels of contrast, the two stimuli were presented 10-15 times during the task. Interleaved measurements of orientation tuning involved 20 presentations of a random sequence of gratings (including the rewarded and unrewarded orientation). Although such a protocol is unlikely to result in SRP, we directly tested this possibility by comparing, in trained animals, evoked potentials across orientations. We computed VEPs traces, separately for six stimulus orientations, by averaging the stimulus-triggered local field potential (LFP) across electrode channels in layer 4 (Figure 10A). We determined VEP amplitudes as the trough to peak difference, and compared them across stimulus orientations (Figure 10B,C). Mean VEP amplitudes did not differ across orientations (main effect of stimulus orientation: p = 0.99, ANOVA), indicating that the learning-related changes are not simply caused by a disproportionate exposure to the rewarded and unrewarded orientation.

**Figure 10.**
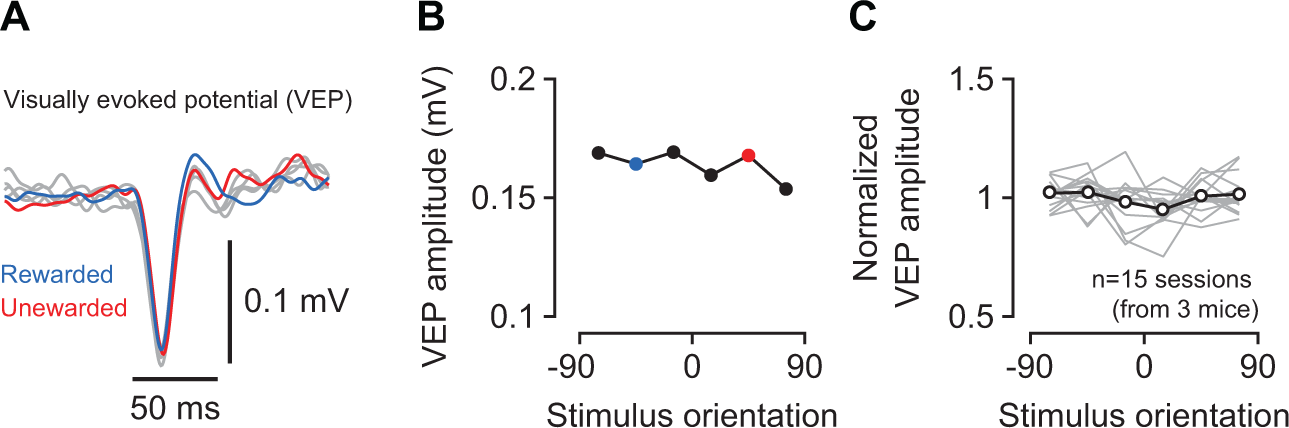
Visually evoked potentials during measurements of orientation tuning. **(A)** VEPs from one recording session in response to gratings of 6 different orientations (M110-36). VEPs for the rewarded and unrewarded stimulus are marked in blue and red. **(B)** Amplitudes of VEPs in (A). **(C)** Normalized VEP amplitude across stimulus orientations for all sessions recorded in the trained stage. Gray lines are individual recording sessions, normalized to the mean amplitude across orientations; black line is the mean across sessions.

## Discussion

We have used a classical conditioning paradigm to investigate how learning about the behavioral relevance of stimulus orientation affects sensory processing in mouse V1. Analyzing behavior in individual animals we found that orientation discrimination learning occurred in a sequence of distinct stages. During an intermediate stage, well before successful discrimination was expressed in the animals’ behavior, we observed substantial modulations of neural activity across the depth of V1: improved discriminability, sharper orientation tuning, and higher contrast sensitivity. We propose that even in a simple learning paradigm, classical conditioning, learning is aided by early modulations of neural responses in visual cortex, which selectively enhance the representation of those sensory signals that are relevant for behavior.

Improvements in sensory processing have long been described in the context of perceptual learning; there are similarities, but also fundamental differences, between this line of research and our study. Perceptual learning refers to improvements in perceptual performance with extensive practice (but see, e.g., Fahle et al., 1995, for faster effects), typically involving threshold-level stimuli (e.g., McKee and Westheimer, 1978; Poggio et al., 1992; Schoups et al., 1995). Our study, in contrast, uses orthogonal stimulus orientations that are far above discrimination thresholds of the mouse (< 10 deg, Andermann et al., 2010; Glickfeld et al., 2013). In perceptual learning sessions, subjects already know which of the stimulus features are relevant to solve the task; in contrast, during initial sessions of our paradigm, the behavioral relevance of grating orientation is to the animal unknown. Rather than investigating neural signatures of perceptual improvements by practice, which can involve thousands of trials (e.g. 150,000 trials, Yang and Maunsell, 2004), we investigated how sensory processing changes as the behavioral relevance of the visual input becomes clear, which could happen after few trials only (see, for instance, Figure 2B). Despite these differences in task requirements and training protocols, there are also important similarities: First, perceptual learning of orientation discrimination can improve selectivity of V1 neurons (e.g., Schoups et al., 2001; Yang and Maunsell, 2004; Raiguel et al., 2006); these improvements, however, were strongest for neurons optimally suited to the task, which contrasts with the global sharpening of tuning we observed. Second, a fraction of our stimuli had contrasts close to threshold, which might have mediated perceptual learning for stimulus contrast. Third, perceptual learning can affect cortical representations outside the context of the task (e.g., Schoups et al., 2001; Yang and Maunsell, 2004; Raiguel et al., 2006; Hua et al., 2010). Fourth, effects of perceptual learning can depend on the level of engagement (e.g., Crist et al., 2001; Yang and Maunsell, 2004; Polley et al., 2006; Li et al., 2008), or the specificity of a task (Li et al., 2004). These similarities between perceptual learning and our approach might indicate that these two forms of learning share fundamental mechanisms of cortical plasticity.

Our finding of distinct changes in behavior is consistent with recent analyses of learning curves in individual subjects. Learning has traditionally been understood as a smooth, hill-climbing process, during which performance gradually increases with the number of trials before reaching an asymptotic level. This process is captured by the classic learning curve, which typically has the form of an inverse exponential function. Such gradually increasing curves, however, might arise from averaging across subjects who, individually, show step-like changes in behavior at different points in time (Gallistel et al., 2004). Indeed, analyses of single-subject data have revealed abrupt changes in a number of learning paradigms, such as autoshaping in pigeons, eye-blink conditioning in rabbits, and proboscis extension in honeybees (Gallistel et al., 2004; Papachristos and Gallistel, 2006; Bazhenov et al., 2013). Similarly abrupt changes are also present in several but not in all of our mice, suggesting that orientation discrimination learning can follow the same dynamics. One way to account for such abrupt changes in behavior is to explain conditioned responses within the framework of perceptual decision-making (Gallistel and Gibbon, 2000; Gallistel et al., 2004).

These sudden changes in behavior, however, do not necessarily imply that learning abruptly changes the neural representation of the stimuli in visual cortex. It might well be that some form of activity-dependent plasticity (Caporale and Dan, 2008) leads to a gradual, trial-by-trial strengthening of synaptic connectivity between neurons along the early visual pathway. Such gradually changing stimulus representations might then be fed to downstream areas, which implement threshold-like decision processes (Bazhenov et al., 2013; Latimer et al., 2015) that finally translate into behavior.

Does learning dynamically shift the preferences of individual V1 neurons? If it did, we should have seen improved discriminability during task performance but not during tuning measurements, potentially in neurons that were driven equally well by the two stimuli (e.g., those with preferred orientations close to 0). Although our data do not suggest any such dynamic shifts of orientation preferences with learning, a decisive answer to this question requires longitudinal tracking of orientation preferences across learning stages. Such chronic tracking during discrimination learning has been achieved with two-photon imaging of mouse V1 (Poort et al., 2015). Although orientation tuning was not measured, these data reveal how the relative response to a rewarded versus unrewarded orientation is affected by learning. In this study, learning reduced day-to-day fluctuations in relative response strength of those neurons that already showed a bias before learning. In addition, learning biased responses of previously indiscriminate neurons towards the rewarded stimulus. Learning has long been know to shift tuning preferences in primary auditory cortex, in particular with fear conditioning (Edeline et al., 1993), but also with other paradigms (Weinberger, 2004). Whether learning can induce comparable shifts in visual cortex requires further research.

The most striking observation in our study is that signatures of learning in V1 were fully expressed well before the behavior indicated that the animals had learned to discriminate the stimuli. The learning effects on V1 responses were at least as strong, sometimes even stronger, in the intermediate than in the trained stage. Such pronounced changes in the spiking responses of V1 neurons early in discrimination learning have not been documented yet. In a number of studies, however, effects of learning have been shown to wax and wane. In primary auditory cortex (A1) of gerbils synaptic inhibition is reduced with learning progress, but returns to pre-training levels when animals become experts (Sarro et al., 2015). Tonotopic representations of sound frequency in rat A1 can expand during initial stages of training and later shrink again (Takahashi et al., 2010), consistent with the observation that cortical map plasticity in A1 is a transient phenomenon occurring during the first few weeks of training (Reed et al., 2011). CA1 neurons in rat hippocampus show enhanced excitability during initial training, but return to pre-training levels with additional practice (Zelcer et al., 2006). Finally, BOLD activity in human V1 increases during learning, but relaxes to pre-training levels with additional training while behavior remains at high levels of performance (Yotsumoto et al., 2008).

Part, but not all, of these early improvements in sensory processing might reflect the presence of a top-down signal, such as attention. We argue that the animals cannot learn this task without attending to the orientation of the stimulus. We speculate that attention is focused in the intermediate stage, and enhances those aspects of the visual scene that are behaviorally relevant. Indeed, the effects of learning we measured are reminiscent of well-known effects of attention on neurons in visual cortex: improved selectivity for stimulus features (David et al., 2008; O’Connell et al., 2014) and improved sensitivity for stimulus contrast (Reynolds et al., 2000; Martinez-Trujillo and Treue, 2002; but see Williford and Maunsell, 2006; Pooresmaeili et al., 2010). Attention is, in fact, a key element in contemporary theories of associative learning (Mackintosh, 1975; Pearce and Hall, 1980; Gottlieb, 2012), and our paradigm might be well suited to reveal attentional effects: low levels of stimulus contrast make anticipation of the consequences difficult, and more difficult tasks typically lead to stronger attentional effects (Chen et al., 2008). Attentional modulation of V1 in human and non-human primates has been shown to change with learning (Gilbert et al., 2000; Bartolucci and Smith, 2011), which might explain the absence of further improvements in the trained stage. Attention, however, cannot be the only factor because improvements in discriminability and sharpening of orientation tuning curves occurred even outside the context of the task.

In addition, these early improvements in sensory processing might reflect a key role of cortex in learning a visually guided task. We hypothesize that during discrimination learning V1 might send a “teaching signal” to subcortical structures, providing information about the visual context, in which the action of anticipatory licking takes place. Such a role of sensory cortex is a key element in a recently proposed model of oculomotor learning (Fee, 2012), which might provide a general framework applicable to a wider range of learning paradigms. From this perspective, V1 would create, already in the intermediate stage, an enhanced representation of the behaviorally important sensory information, which is used to learn the mapping between stimuli and outcomes.

## Conflict of interest

The authors declare no competing financial interests.

## Acknowledgements

We thank Daniel Fürth for valuable advice, and Dimitri Yatsenko & Alexander Ecker for sharing code for data management (DataJoint, http://datajoint.github.io/). This work was supported by a Starting Independent Researcher grant from the European Research Council awarded to S.K. (281885 PERCEPT) and by funds awarded to the Centre for Integrative Neuroscience within the framework of the German Excellence Initiative (DFG EXC 307).

